# Timing of gene expression in a cell-fate decision system

**DOI:** 10.1101/197103

**Authors:** Delphine Aymoz, Carme Solé, Jean-Jerrold Pierre, Marta Schmitt, Eulàlia de Nadal, Francesc Posas, Serge Pelet

**Affiliations:** Department of Fundamental Microbiology, University of Lausanne, Lausanne, Switzerland; Cell Signaling Research Group, Departament de Ciències Experimentals i de la Salut Universitat Pompeu Fabra, Barcelona Spain

## Abstract

During development, morphogens provide extracellular cues allowing cells to select a specific fate by inducing complex transcriptional programs. The mating pathway in budding yeast offers simplified settings to understand this process. Pheromone secreted by the mating partner triggers the activity of a MAPK pathway, which results in the expression of hundreds of genes. Using a dynamic expression reporter, we quantified the kinetics of gene expression in single cells upon exogenous pheromone stimulation and in the physiological context of mating. In both conditions, we observed striking differences in the timing of induction of mating-responsive promoters. Biochemical analyses and generation of synthetic promoter variants demonstrated how the interplay between transcription factor binding and nucleosomes contribute to determine the kinetics of transcription in a simplified cell-fate decision system.

**One Sentence Summary:** Quantitative and dynamic single cell measurements in the yeast mating pathway uncover a complex temporal orchestration of gene expression events.

## Main Text

Cell-fate decisions play a key role in embryonic development. In order to make choices, cells integrate cues from neighboring cells as well as from morphogens. Signal transduction cascades relay this information inside the cell to translate these extra-cellular signals into defined biological responses. The cellular output includes the induction of complex transcriptional programs where specific genes are expressed to different levels and at various times (*1, 2*). Ultimately, these different expression programs will determine the fate of individual cells. The mating pathway in budding yeast has often been considered as a simplified cell-fate decision system, where each cell can either continue to cycle in the haploid state or decide to mate with a neighboring cell of opposing mating type. This decision results in an arrest of the cell cycle, formation of a mating projection and ultimately leads to the fusion with the partner to form a diploid zygote (*3, 4*).

Haploid budding yeast senses the presence of potential mating partners by detecting pheromone in the medium. This small peptide elicits the activation of a Mitogen-Activated Protein Kinase (MAPK) cascade (Sup Fig 1), which can integrate multiple cues such as stresses, cell cycle-stage or nutrient inputs (*5-8*). Once the MAPKs Fus3 and Kss1 are activated, they phosphorylate a large number of substrates and induce a new transcriptional program. Ste12 is the major transcription factor (TF) implicated in this response, and controls the induction of more than 200 genes (*9*). Under normal growth conditions, this TF is repressed by Dig1 and Dig2. Phosphorylation by active Fus3 and Kss1 relieves this inhibition, such that Ste12 can recruit the transcriptional machinery (*10*). Ste12 associates to the DNA via well-established binding sites located in promoters called Pheromone Response Elements (PRE), with the consensus sequence ATGAAACA (*11, 12*). Although PREs are found upstream of the vast majority of pheromone-induced genes (*13*), the number of binding sites, their orientation and their position relative to the transcription start site vary widely from one gene to the next (*13, 14*).

**Figure 1.**
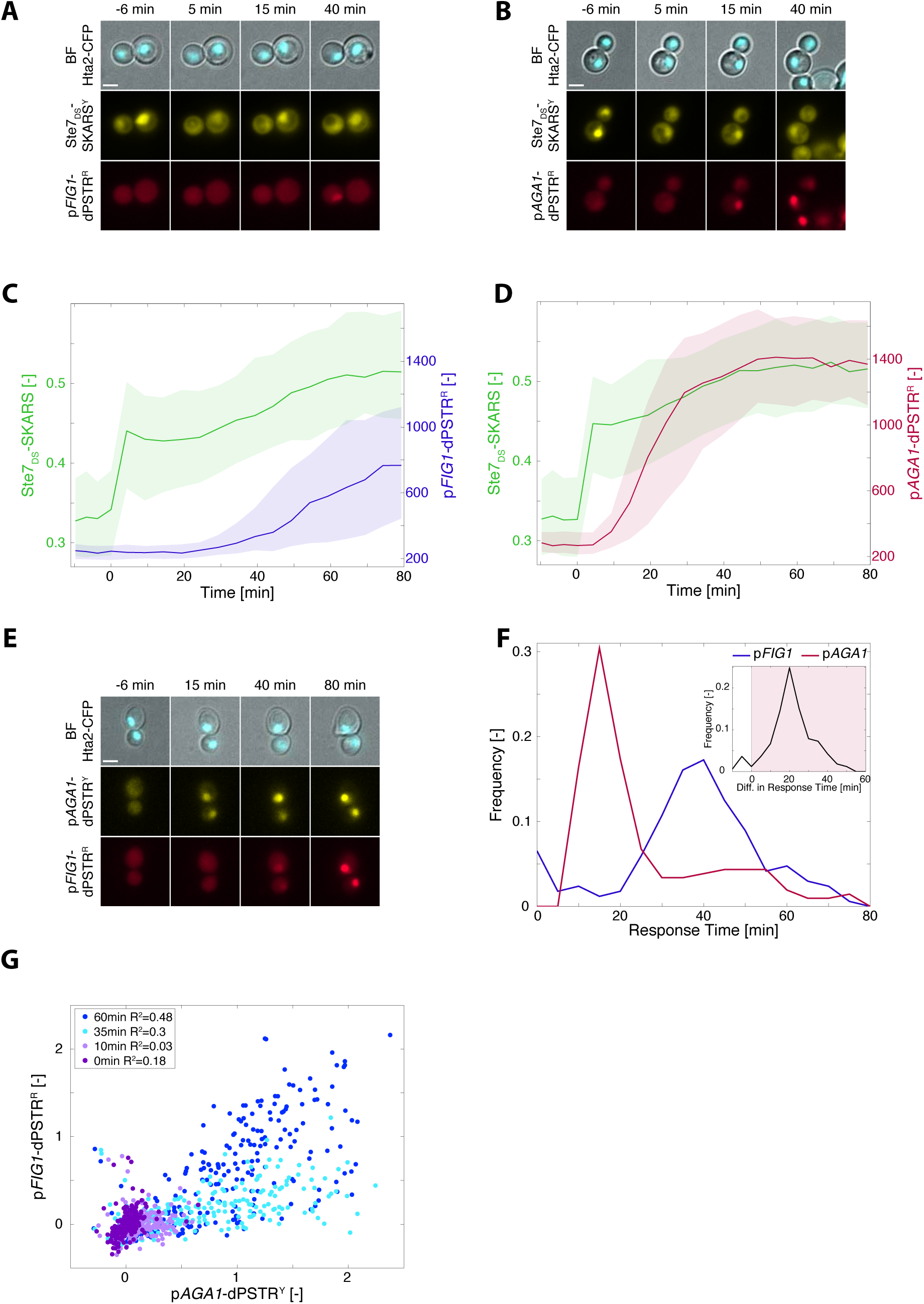
Interplay between kinase activity and promoter induction in the mating pathway. **A and B.** Microscopy images of cells stimulated with a saturating pheromone concentration (1 µM) at time 0 minutes. The cells bear a histone tagged with CFP, a yellow SKARS reporting on Fus3p and Kss1p activities, and a red dPSTR reporting on p*FIG1* (A) or p*AGA1* (B) induction. For all experiments, unless stated otherwise, the stimulation was performed by addition of 1 µM α-factor at time 0 minutes. All scale bars on microscopy images represent 2.5 µm. **C and D.** Quantifications of the kinase activity (green, left axis), measured by the ratio of cytoplasmic to nuclear YFP, and of the p*FIG1* (C) and p*AGA1* (D) expressions, measured by the difference between nuclear and cytoplasmic fluorescence of the dPSTR (right axis). For all similar graphs, the solid line is the median response and the shaded area represents the 25 to 75 percentiles of the population. **E.** Microscopy images of a strain carrying p*FIG1*-dPSTR^R^ and p*AGA1*-dPSTR^Y^. **F.** Quantification of the response time of p*FIG1* and p*AGA1* reporters (see Methods). The inset is the difference response time between the p*AGA1*-dPSTR^Y^ and the p*FIG1*-dPSTR^R^, for all cells expressing both promoters. The red shaded area represents cells expressing p*AGA1* before p*FIG1* (87%). **G.** Correlation of normalized dPSTR nuclear enrichments from all single cells of a representative experiment at different time points after stimulation.

Promoter sequences are primary determinants of the strength and kinetics of gene expression. Unfortunately, the basic rules governing transcription regulation remain poorly understood. Libraries of synthetic promoter sequences have allowed establishing a few rules in the control of the expression level and the noise of a promoter sequence (*15-17*). However, the slow maturation time of fluorescent proteins (FP) precluded thorough investigations of gene expression kinetics.

In this study, we used dynamic gene expression reporters (*18*) to characterize the expression dynamics of a set of promoters induced in response to yeast mating pheromone. We have identified different classes of promoters based on the kinetics of their expression. Deeper analysis of early and late promoters highlighted the interplay between TF binding and nucleosome positioning as a major determinant of the expression dynamics. In addition, we demonstrate that under physiological mating conditions, the induction of the target genes follows a precise chronology, and are sequentially expressed until fusion occurs.

### Interplay between kinase activity and expression dynamics

In multiple MAPK pathways, MAPK activity has been shown to be tightly linked to the transcriptional process by phosphorylating TFs, contributing to the recruitment of remodeling complexes, and participating in the elongation complex (*19*). Therefore, we wanted to measure, in the mating pathway, how kinase activity and gene expression were temporally correlated. Using fluorescent relocation sensors that we previously engineered, we are able to quantify, in real-time and at the single cell level, both MAPK activity and gene expression upon stimulation of *MATa* cells with synthetic pheromone (α-factor, 1*µ*M) (*18, 20*). Signaling activity was quantified using a Ste7_DS_-SKARS^Y^, which exits the nucleus when the mating MAPKs Fus3 and Kss1 phosphorylate specific residues in the vicinity of a nuclear localization sequence (NLS) (Fig 1A; Sup Fig 2A). In the same cells, a dynamic protein expression reporter p*FIG1*-dPSTR^R^ was integrated. *FIG1* displays the largest fold-induction upon pheromone stimulation (*9*). In this assay, the *FIG1* promoter drives the expression of a small peptide, which interacts with a fluorescent protein and promotes its recruitment in the nucleus (Fig 1A, Sup Fig 2B, (*18*)). Upon stimulation, the cells activate the mating MAPKs few minutes after stimulation, as previously described (*5, 20, 21*). Despite this fast signal transduction, the resulting p*FIG1* expression occurs 30 minutes later (Fig 1A and C). Individual yeast cells are known to possess a large diversity in signaling capacity (*6, 22*). Analysis of the sub-population that activates strongly the MAPK within the first 10 minutes after stimulus still results in a delayed and broad distribution in p*FIG1* activation dynamics (Sup Fig 3A). This finding suggests an absence of temporal correlation between kinase activity and the downstream transcriptional response.

**Figure 2.**
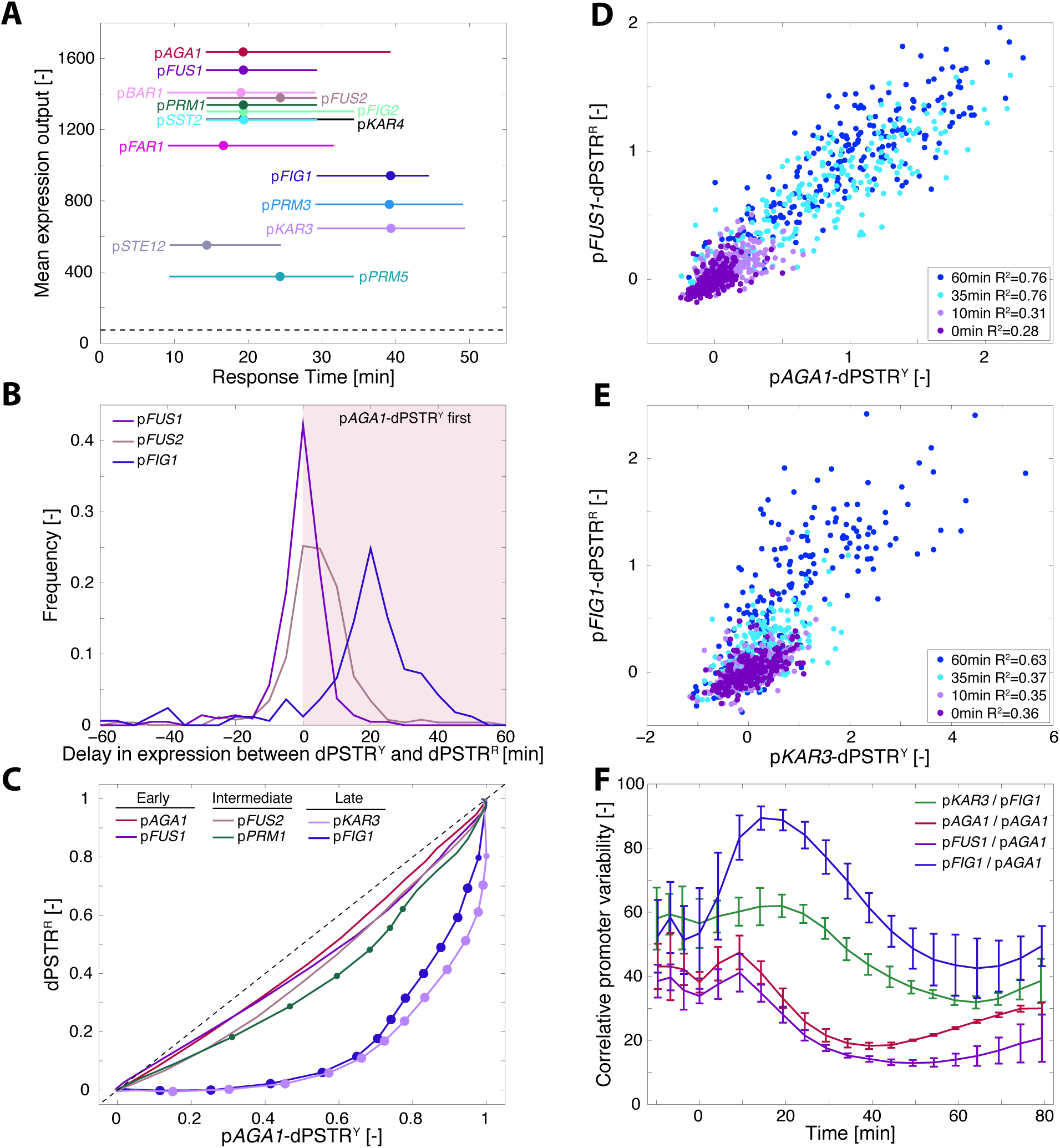
Dynamics of induction of mating promoters after pheromone stimulation. **A.** Response time versus mean expression output for the 14 mating-dependent promoters. Dots represent the median response times of the cell population and lines represent the 25^th^ and 75^th^ percentiles. All promoters were measured with the dPSTR^R^. The strains also bear the p*AGA1-*dPSTR^Y^ for direct comparison of the dynamics of promoter induction. The dashed line represents the detection sensitivity of the dPSTR^R^ reporter. **B.** Distributions of the differences in the response times between the p*AGA1*-dPSTR^Y^ and the dPSTR^R^ in the same cell for p*FUS1*, p*FUS2* and p*FIG1*. **C.** Correlation of the population averaged normalized nuclear enrichment of p*AGA1*-dPSTR^Y^ and a selected set of promoters measured with the dPSTR^R^ at all time points of the experiments. The dots represent the p-Value (10^−3^>pval>10^−6^ for small dots and pval <10^−6^ for large dots) of the *t*-test comparing the offset of the measured promoter relative to the x=y line with the offset of the reference promoter p*AGA1*. **D and E.** Correlation of normalized dPSTR nuclear enrichments of single cells of at different time points after stimulation in a strain with p*FUS1*-dPSTR^R^ and p*AGA1*-dPSTR^Y^ (D) or p*FIG1-*dPSTR^R^ and p*KAR3*-dPSTR^Y^ (E). **F.** Evolution of the Correlative Promoter Variability (CPV) in course of time, for various pairs of promoters. The curve represents the mean of 3 replicates and the error bar the standard deviation between replicates. A low CPV corresponds to a similar expression between two promoters in the same cell (see Methods).

This surprising result led us to test the expression kinetics of multiple mating responsive promoters. Among them was *AGA1*, a gene reported to be strongly induced upon pheromone stimulation (*9, 23*). The p*AGA1*-dPSTR^R^ begins to enrich in the nucleus of cells 15 minutes after stimulation (Figure 1B and D). Thus, the induction of gene expression from this promoter is much faster than for p*FIG1*. In addition, the induction of p*AGA1* in signaling competent cells is less variable with the vast majority of the cells inducing the reporter within the 30 minutes following the stimulus (Sup Fig 3A). This raises the question of how the expression of these two promoters is related in a same cell.

**Figure 3.**
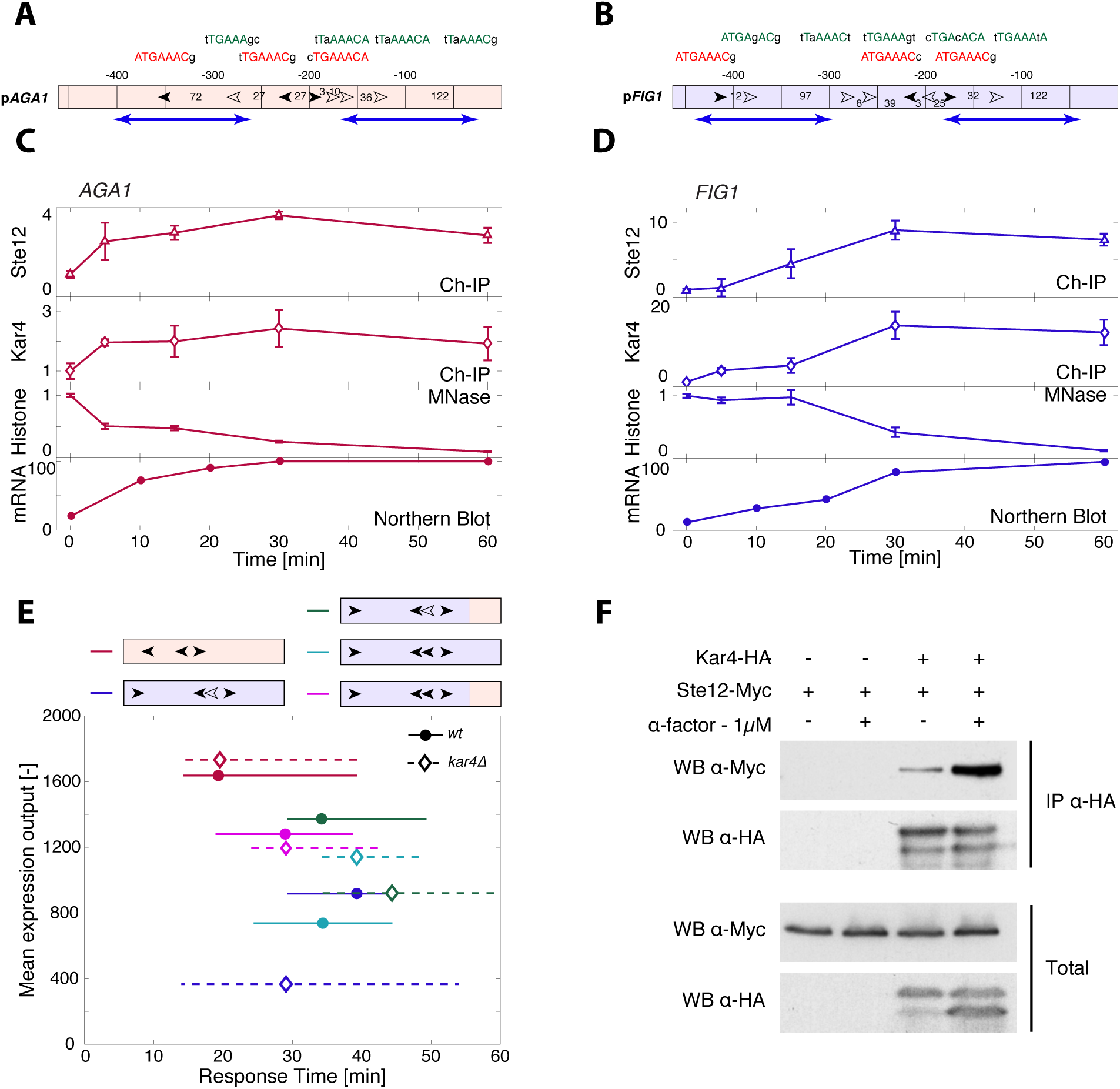
Influence of promoter architecture on expression dynamics. **A and B.** Maps of the two promoters p*AGA1* and p*FIG1.* The filled arrows represent the location and orientation of consensus Ste12-binding sites (nTGAAACn). The open arrows symbolize the non-consensus binding sites, that possess mutations within the 6 core nucleotides of the PREs. The sequences of each binding sites is detailed above, with capital nucleotides matching the consensus sequences and small nucleotides being mutations from the consensus. The numbers between sites represent the distance in bp between them or the ATG. Blue arrows represent nucleosomes position (*27*). **C and D.** Quantification of molecular events at the AGA1 (C) and FIG1 (D) loci. Fold increase in Ste12-myc and Kar4-HA binding at the promoter quantified by chromatin-IP (open markers). Normalized −1 histone occupancy quantified by Micrococcal Nuclease (MNase) digestion. Transcript levels of *AGA1* (C) and *FIG1* (D) quantified by Northern Blot (rounds). **E.** Response time versus mean expression output for various promoters in a WT background (circles, solid lines) or *kar4Δ* (diamonds, dashed lines) background, as described in Figure 2A. Red is p*AGA1*, blue is p*FIG1*, green is a chimeric construct between p*FIG1* and the last 150 bp of p*AGA1*, cyan is a construct where the free non-consensus binding site of p*FIG1* (-209) was mutated into a consensus one, and purple is a combination of the chimeric construct with the mutation of the non-consensus binding site into a PRE. **F.** *In vivo* binding of Ste12 and Kar4 was assessed by immunoprecipitation of Kar4p-HA and detection of Ste12-Myc in presence and absence of pheromone.

### Direct comparison of two dynamic expression reporters

We used a second protein expression reporter, the dPSTR^Y^, which is orthogonal to the dPSTR^R^, to quantify p*AGA1* and p*FIG1* expression dynamics in the same strain (*18*) (Figure 1E and Sup Fig 3B). In all expressing cells, the response time for each promoter was determined based on the time at which the dPSTR nuclear enrichment reached 20% of its maximum (Figure 1F, see methods and Sup Fig 4). p*AGA1* expression is relatively homogeneous between cells, with 83% of the cells inducing the promoter within the first 30 minutes following stimulation. In comparison, p*FIG1* expression is highly variable from cell to cell. In cells inducing both promoters, the difference in response times can be measured (Figure 1F, inset). In 87% of cells, the p*AGA1*-dPTSR^Y^ is activated prior to the p*FIG1*-dPSTR^R^,which on average is delayed by 23 minutes. These different dynamics of induction are also well illustrated by the absence of correlation between the dPSTR enrichment seen at early time points (Figure 1G). The cell population becomes first p*AGA1* expressing, as denoted by a shift along the *x*-axis. Then, the population moves upwards, illustrating the delay in the induction of p*FIG1*. This delay is not an artifact from the dPSTRs, since the same results can be obtained when exchanging the promoters on the dPSTRs (Sup Fig 5 and 6). In parallel, we have also verified that mRNA production dynamics from these two promoters correlate well with the expression dynamics we measured with the dPSTR (Sup Fig 7). Together, these data demonstrate that although the MAPK activity rises quickly in response to pheromone sensing, it does not lead to a fast and simultaneous activation of all mating genes.

**Figure 4:**
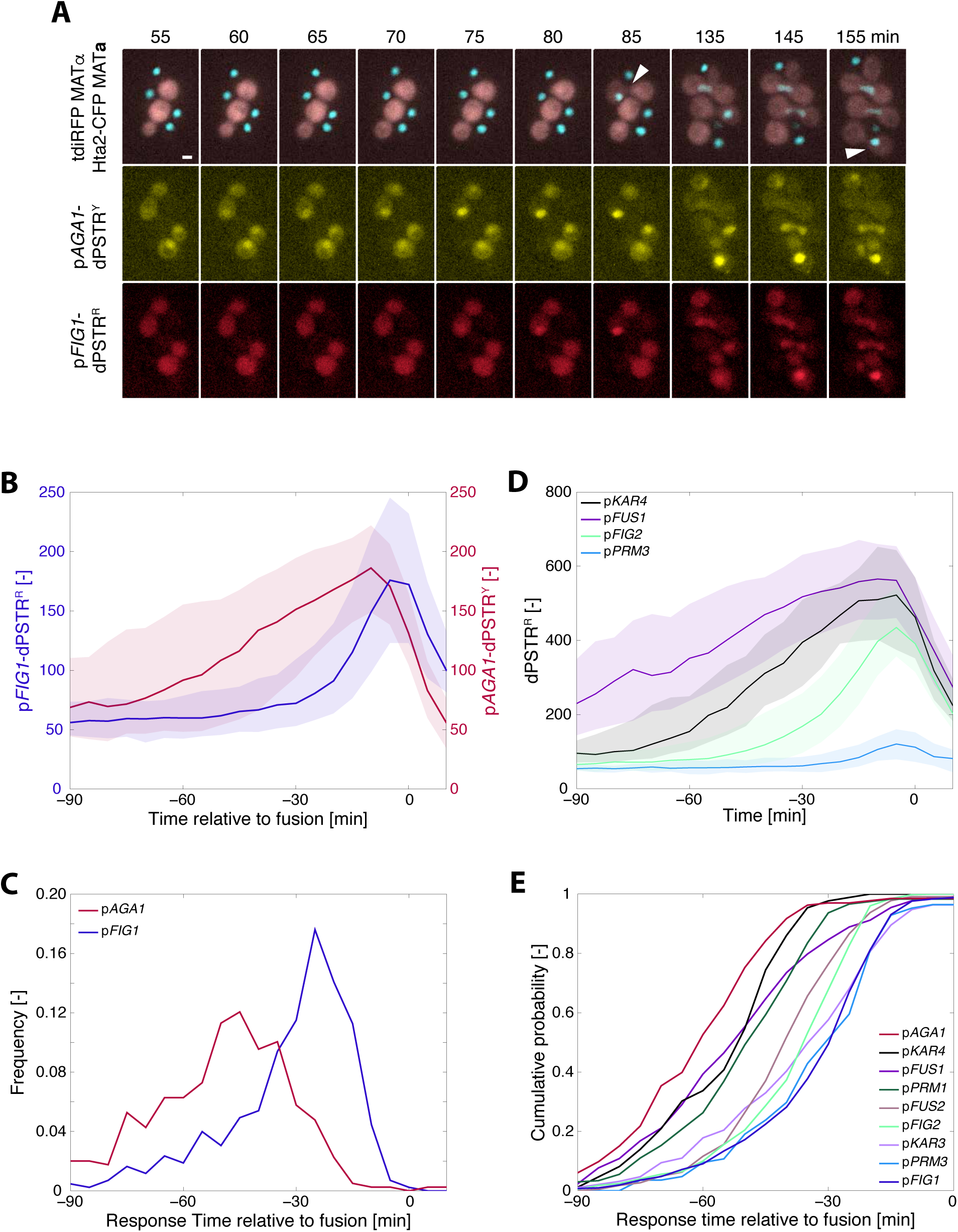
Dynamics of gene expression during the mating process. **A.** Microscopy images of a mating mixture containing the MATa strain (Hta2-CFP, p*FIG1-*dPSTR^R^ and p*AGA1*-dPSTR^Y^) and a MATα (cytoplasmic tdiRFP) at different times after beginning of the imaging (time 0). Fusion events are marked by a white arrow. **B.** Quantification of the nuclear enrichment of p*FIG1*-dPSTR^R^ (blue, left axis) and of p*AGA1-*dPSTR^Y^ (red, right axis). Single cell traces were synchronized relative to their fusion time, identified by a sudden increase in tdiRFP signal into the MATa cells. **C.** Distribution of the response time of p*AGA1* and p*FIG1* relative to the fusion time. **D.** Activation dynamics of various promoters prior to fusion as measured by dPSTR^R^ in different mating mixtures. **E.** Cumulative probability of the response time relative to fusion for 9 mating induced promoters measured in mating conditions.

### Characterization of mating-induced promoters

Having established that the two promoters p*AGA1* and p*FIG1* are expressed with different kinetics following pheromone stimulation, we tested when other mating-induced genes were expressed with respect to p*AGA1*. Fourteen mating-responsive promoters, previously described in the literature, were characterized using a dPSTR^R^ (Sup Fig 8) (*9, 13, 14*). We quantified for each of them their expression output, by measuring the variation of the nuclear enrichment of the dPSTR^R^ upon stimulation (see methods, Sup Fig 4). These promoters display a large variability both in the level of induction and the timing of expression (Figure 2A). Some genes are expressed early as *AGA1* (*FUS1, FAR1, STE12*, etc…), others are late responders similar to *FIG1* (*PRM3, KAR3*).

In order to better characterize the dynamics of expression of the 14 reporters, they were compared to the same internal control, a p*AGA1*-dPSTR^Y^. The difference in response time relative to p*AGA1* induction was calculated (Figure 2B, Sup Fig 9). In addition, the comparison of the overall dynamics of induction was visualized by plotting the mean nuclear enrichment of the yellow and red dPSTRs, normalized between their basal and maximal expression levels (Fig 2C, Sup Fig 9). Each curve starts in the lower left corner and evolves towards upper right corner as promoters are being induced during the time-lapse. Promoters which are induced with similar dynamics as p*AGA1*, will remain close to the *x*=*y* diagonal (dashed line). Any difference in induction dynamics will deviate from this line. Based on these measurements, we defined three classes of promoters: early, intermediate and late. The early promoters, with kinetics similar to p*AGA1,* display a difference in response time centered around zero and a correlation aligned on the *x*=*y* diagonal (Sup Fig 9). Late promoters, which behave similarly to p*FIG1*, have a response time delayed by at least 15 minutes and a correlation strongly deviating from the diagonal. Between these two clearly identifiable groups, a set of promoters display intermediate kinetics, where the response time is slightly delayed and/or where the dynamic correlation with p*AGA1* is significantly deviating from the p*AGA1*/p*AGA1* correlation at many time points.

The basal level of expression before stimulus (Sup Fig 10) or the maximal expression level reached after pheromone induction (Fig 2A) do not allow to predict whether a promoter will be fast or slow. For instance, the *STE12* promoter belongs to the early genes group, but possesses one of the lowest induction levels. However, there is a clear link between the ability to respond at low pheromone concentration and the dynamics of promoter induction (Sup Fig 11 and 12). p*AGA1* and other promoters from this category display a graded response as α-factor concentration increases. In comparison, late promoters behave in a more switch-like manner, where gene expression occurs only at high concentrations of α-factor (300 nM).

### Variability in gene expression

When focusing on the single cell responses, a remarkable correlation between the expression of the fast promoters at various time points can be observed (p*AGA1*/p*FUS1*: Fig 2D and other pairs in Sup Fig 13 and 14). This tight correlation can be explained by the low noise present in the mating pathway and the expression variability being mostly governed by extrinsic variables such as the cell cycle stage and the expression capacity (*22*). More striking is the fact that two late promoters in the same cell are also induced with a good correlation. This implies that despite the fact that the induction of these late genes can occur 30 to 80 minutes after the stimulus, these two promoters are activated synchronously within a given cell (Fig 2E and Sup Fig 6 and 14). These data also allow to rule out the presence of a slow stochastic activation of the late genes and rather argue in favor of a specific commitment point that the cells reach when they start to induce the late promoters.

In order to illustrate this better, we defined the Correlative Promoter Variability (CPV), which allows to quantify the deviation in the induction of two promoters measured in the same cell, relative to the overall noise in expression. (Fig 2F and Sup Fig 6 and 9, see methods). For two promoters well correlated like p*AGA1* and p*FUS1* (*24*), the CPV starts below 50%, and tends to further decrease upon pheromone-dependent induction. Among fast promoters, there can be different types of behavior, depending mostly on the pre-stimulus levels of the reporter. The variability between p*FAR1* and p*AGA1* is a good illustration of this (Sup Fig 9). The CPV is high in basal conditions, due to the asynchronous induction of p*AGA1* and p*FAR1* during the cell cycle (Sup Fig 10 and 13, (*24*)). However, following stimulation with pheromone, the variability decreases quickly as the two promoters are simultaneously induced. In comparison, the CPV between the late *FIG1* promoter and the early p*AGA1* increases during the first 20 minutes following induction, due to an asynchronous induction of p*AGA1* and p*FIG1*. Upon activation of the late promoter, the variability decreases. For the two slow promoters p*FIG1* and p*KAR3*, the basal CPV value is around 60%, due to uneven basal levels of the p*KAR3* induction during the cell cycle ((*25*), Sup Fig 10). After stimulus, this CPV level is maintained for roughly 30 minutes, during which none of these two promoters are induced and then drops. Overall, these measurements demonstrate that each mating-induced promoter is expressed with specific dynamics and expression level. Some cells will induce the early genes few minutes after the stimulus, while late gene expression can be delayed by more than an hour. Remarkably, the tight co-regulation of early and late genes within their group strongly suggests that a shared mechanism exists that regulates the early promoters, which is different from the one controlling the activation of the late promoters.

### Architecture of mating promoters

In order to understand how the timing of induction is regulated, we have mapped all putative Ste12 binding sites in the sequences of the fourteen promoters (Sup Fig 15). We defined consensus PREs as nTGAAACn, as it was reported that these six core nucleotides were the most important to promote Ste12 binding *in vitro* (*14*). We also identified several non-consensus PREs that carry additional mutations within the six core nucleotides. These putative binding sites possess a decreased affinity for Ste12, but can contribute to Ste12-mediated expression (*14*). As reported previously, there is a large variability in the number, orientation, spacing and sequences of PREs among all promoters (*13, 14*). Therefore, there is no obvious rule that would allow to predict whether a gene is early-or late-induced, or expressed at low or high levels. Interestingly, p*AGA1* and p*FIG1* possess 3 consensus PREs with relatively similar dispositions and orientations, and respectively four and five non-consensus PREs (Fig 3A and B). Despite these similarities, we have observed drastic differences in their expression kinetics. Therefore, we decided to use p*AGA1* and p*FIG1* as model promoters for their categories and decipher their mode of regulation.

### Regulation of pAGA1 and pFIG1

In a strain bearing the p*FIG1*-dPSTR^R^ and the p*AGA1*-dPSTR^Y^ reporters, key regulators of the pathway were deleted. A number of mutants did not affect the expression from both promoters (group I: *kss1Δ, tec1Δ mot3Δ* and *arp8Δ* Sup Fig 16) or altered it in a similar fashion (group II: *ste12Δ, ste2Δ, ste11Δ, dig1Δdig2Δ*, Sup Fig 17). However, the interesting knockouts are the ones that perturbed one promoter to a greater extent than the other one (group III). In *fus3Δ* and *far1Δ* cells, p*AGA1* induction is delayed while p*FIG1* is severely reduced. Only a small percentage of cells induce p*FIG1* (Sup Fig 18). Cells deleted for a member of the SAGA chromatin remodeling complex (*gcn5Δ*) also displayed a stronger decrease in p*FIG1* induction than in p*AGA1* suggesting a higher requirement for nucleosome remodeling at the *FIG1* than at the *AGA1* promoter. Finally, deletion of the transcription factor *KAR4* profoundly affects p*FIG1* induction, without noticeable changes in p*AGA1*-dPSTR^Y^ expression. Kar4 has been identified as a transcription factor required for the induction of genes implicated in karyogamy, a late event of the mating (*25*). Microarray measurements have identified a set of genes, such as *KAR3* and *PRM3,* to be dependent on Kar4 but not *FIG1* (*26*). It has also been suggested that Kar4 forms a heterodimer with Ste12 and therefore the association of those two proteins on the promoter allows the transcription of the late genes (*26*). Moreover, we found that *KAR4* is induced as early as p*AGA1* during the mating response, making it a good candidate to regulate late genes.

### Ste12 and Kar4 interplay at the promoter

In order to better understand the sequence of events taking place at these two promoters, we monitored transcription factor binding by chromatin-IP, chromatin remodeling by MNase assays and mRNA production by Northern Blot. All these experiments were performed in the same strain with Ste12-myc and Kar4-HA tags. We noticed that the presence of these tags slightly influences the dynamics of transcription although the differential response of the two promoters is maintained (Sup Fig 19 A and B). On the *AGA1* promoter, a fast enrichment of Ste12 and Kar4 is observed within 5 minutes after stimulus. In parallel, the chromatin is remodeled on the locus, as visualized by the eviction of the −1 histone (Fig 3C and Sup Fig 19C). The concomitant enrichment in TF and opening of the chromatin results in a rapid production of mRNA. In comparison, at the *FIG1* locus all these events happen more slowly (Fig 3D and Sup Fig 19D). Ste12 seems to accumulate first, followed by Kar4, and chromatin remodeling takes place later, around 30 minutes after the stimulus. As a consequence, the resulting mRNA production is delayed at this locus.

The ability of TFs to bind promoter regions is known to depend on the positioning of nucleosomes on the DNA. MNase protection assays, in agreement with genome-wide studies (Sup Fig 19 C and D (*27*)) allow to predict which PRE could be accessible under basal conditions. On p*AGA1*, two consensus binding sites for Ste12 are present in a nucleosome-depleted region (Fig 3A). This conformation would allow the formation of a Ste12 dimer under basal conditions. Indeed, both Ste12 and Kar4 are found associated with *AGA1* and *FIG1* promoters even before the addition of a-factor (Sup Fig 19E). The Ste12 dimer on p*AGA1* could allow a fast induction of transcription as soon as Fus3 activity is present to derepress Dig1 and Dig2. In agreement with this prediction, mutation of either of these PRE sites delays significantly the induction of p*AGA1* transcription, and mutation of both PREs virtually abolishes the induction of this promoter variant (Sup Fig 20A to C).

In comparison, only one strong Ste12 binding site is found in a nucleosome-depleted region of p*FIG1* (Fig 3B). This site lies in the close vicinity of a non-consensus site. Surprisingly, mutation of either of these two sites completely abolishes the mating-dependent induction from these promoter variants (Sup Fig 20D and E). In order to understand the parameters that control the dynamics of induction of the late promoters, we performed a series of mutations to test if we achieved to accelerate the dynamics of induction of the p*FIG1* promoter. In a first variant, we mutated the non-consensus site of p*FIG1* into a consensus one. This operation could putatively allow the recruitment of a Ste12 dimer under basal conditions, because both binding sites fall in a nucleosome-depleted region of the *FIG1* locus. This promoter variant turned out to be only marginally faster than the WT promoter. However, this single point mutation in the non-consensus site renders the induction of this promoter Kar4-independent (Figure 3E, Sup Fig 20 F and G).

To alter the nucleosome landscape on p*FIG1*, we constructed a promoter chimera and replaced the 150 bp of the core promoter that are associated with −1 nucleosome in p*FIG1* by the p*AGA1* sequence. This promoter chimera displays an intermediate behavior between p*FIG1* and p*AGA1*. It is faster and more expressed than the natural p*FIG1* promoter and retains a Kar4 dependency. By combining these two modifications (non-consensus to consensus PRE in the chimera) we further accelerated the induction of the dPSTR and rendered it Kar4-independent (Fig 3E).

Taken together, these data allow us to infer a model where early genes possess at least two consensus binding sites for Ste12 in a nucleosome-depleted region, an assumption true for all fast promoters tested in this study except p*PRM1*. Activation occurs rapidly via the inhibition of Dig1/2 in a manner that is proportional to the pheromone concentration and signaling activity present in the cell. Late genes do not have the ability to form these Ste12 dimers under basal conditions, because at most one consensus Ste12 site is found in a nucleosome-depleted region. Based on the evidences provided here, we postulate that the formation of a Ste12 dimer using non-consensus sites can be stabilized by Kar4. Interestingly, Kar4 has been found associated to the *AGA1* promoter in basal condition, but its deletion does not alter the level of expression or the dynamics of induction of this early promoter. However, the dynamics of induction of intermediate promoters are perturbed in a *kar4Δ* background (Sup Fig 21). Therefore, our data demonstrate a more global effect of Kar4 on mating genes induction than previously thought. We also observed an interaction between Ste12 and Kar4 that is strongly enhanced by pheromone treatment (Fig 3F). The association between Ste12 and Kar4 is needed to recruit Kar4 on the promoter, as in *ste12Δ* cells Kar4 is not detected on p*AGA1* or p*FIG1* (Sup Fig 19E and F). Kar4 presence could stabilize the TF complex on the promoter allowing a recruitment of the chromatin remodelers, so as to evict the nucleosomes and induce an efficient transcription of the downstream ORF. The delay observed in the late genes expression is thus a combination of the requirement for Kar4 to be transcribed at sufficient levels to allow interaction with Ste12 and slow chromatin remodeling on these loci. Both Ste12-Kar4 interaction and chromatin remodeling are enhanced by MAPK activity (*28*), which can explain the requirement for a high pheromone concentration, and thus an elevated kinase activity, to induce the late promoters.

### Promoter induction during mating

The characterization of the various mating-dependent promoters has been performed in well-controlled conditions using synthetic mating pheromone. We next wanted to verify whether similar dynamics of gene expression occurred under the physiological conditions of mating. *MATa* cells bearing the p*FIG1* and the p*AGA1* dPSTRs were mixed on an agar pad with *MAT*α cells constitutively expressing an infrared FP (tdiRFP) (Fig 4A). Strikingly, under these conditions, we also observed a clear difference in the activation of the two reporters. The *AGA1* promoter is already induced in some cells at the onset of the time-lapse (∼30 minutes after the mixing of the mating partners). As time goes by, more cells induce the *AGA1* reporter (Fig 4A). In comparison, the p*FIG1*-dPSTR^R^ is expressed in fewer cells and its induction precedes the fusion of the partners. Using an automated image analysis pipeline, fusion events can be detected in *MATa* cells by a strong and sudden increase in tdiRFP fluorescence (Sup Fig 22). The single cell traces of 455 of these events recorded in one experiment were aligned temporally to their fusion time, set to 0. These quantifications reveal very clearly that the induction of p*AGA1* gradually increases until it reaches a peak prior to fusion (Fig 4B). In comparison, the *FIG1* promoter is not active until roughly 30 minutes before fusion. The measurements of the response time relative to fusion confirm the kinetic difference between p*AGA1* and p*FIG1.* In addition, these new findings indicate that p*FIG1* induction seems to be tightly correlated with the fusion time, while p*AGA1* is expressed earlier and with a larger variability (Fig 4C). Cells that did not undergo fusion are highly likely to induce p*AGA1*, while p*FIG1* induction is rare in this sub-population. It can be sometimes observed in cells in the close vicinity of a set of engaged mating partners (Sup Fig 23).

We verified that this difference in dynamics of expression is also present for other promoters (Fig 4 D and E, Sup Fig 24). In agreement with our classification based on exogenous stimulations experiments, early genes are the first ones to be induced in the mating process, followed closely by intermediate genes. Late genes induction precedes the fusion time by only 30 minutes, a time when cells seem committed to this process. Therefore, these genes are rarely being expressed in non-fusing cells, which is not the case for early and intermediate genes.

These experiments provide a better understanding of the key steps in the mating process. As soon as mating pairs are in proximity, the low level of pheromone constantly produced by the cells is sufficient to trigger an activation of the mating pathway and induction of the expression of early mating genes. Many of these early genes are implicated in sensing and cell-fate determination, and will contribute to the commitment of the partners to the mating process. If both partners are able to arrest in G1, they will extend a mating projection towards each other and polarize their sensing and secretory machinery. This will lead to a local increase in pheromone concentration that will be associated with an increase in signal transduction (Sup Fig 25 and (*29*)). Mating experiments performed with a mutant unable to degrade pheromone (*bar1Δ*) clearly demonstrate that p*FIG1* induction is triggered by the concentration of pheromone sensed by the cells and not by cell-cell contacts. In this mutant, non-fusing cells activate this promoter because they experience a high concentration of pheromone independently of their proximity to a mating partner (Sup Fig 26). Taken together, our results demonstrate that yeast cells use a temporal gradient of pheromone to orchestrate the timing of expression of mating genes.

This behavior bears many similarities with morphogen sensing in development. Concentration of the diffusive signal was thought to be the key element for cell-fate decision. It is now apparent that both level and timing of morphogen stimulus dictate early and late gene expression (*1, 30-33*). A key question is how this temporal information is encoded to deliver the proper gene expression profile. In the simple settings offered by budding yeast, our data show that both the affinity of the TF binding sites and chromatin state at the promoter determine the concentration threshold and the timing of gene expression. This may be a general mechanism of how the timing of gene induction is orchestrated in a wide variety of cell-fate decision systems.

## Acknowledgments

We thank all members of the Pelet and Martin labs for helpful discussions and suggestions. We thank Sophie Martin and Laura Merlini for critical readings of the manuscript. We are grateful to Clémence Varidel, Réjane Seiler, Joan Jordan and Christine Boaron for technical assistance. Work in the Pelet lab is supported by the Swiss National Science Foundation, SystemsX.ch and the University of Lausanne. Work in the Posas and de Nadal laboratories is funded by grants from the Spanish Ministry of Economy and Competitiveness (BFU2015-64437-P and FEDER, BFU2014-52125-REDT, and BFU2014-51672-REDC to FP; BFU2014-52333-P and FEDER to EN), the Catalan Government (2014 SGR 599), and the FundaciÓn Botín, by Banco Santander through its Santander Universities Global Division to FP. FP is recipients of an ICREA Acadèmia (Generalitat de Catalunya).

## Authors’ contributions

SP and DA conceived the study. SP, DA, FP and EN designed the experiments. MS and JJP built plasmids and strains. DA, SP and JJP performed the microscopy experiments and analyzed the data. CS performed biochemistry experiments. SP and DA wrote the paper. All authors read and approved the final manuscript.

## Supplementary Materials

Materials and Methods

Figs. S1 to S26

Tables S1 to S3

